## Supplementary Information for "msiPL: Non-linear Manifold and Peak Learning of Mass Spectrometry Imaging Data Using Artificial Neural Networks"

### Supplementary Materials and Methods:

MSI datasets from five different tissue types were analyzed using the neural network model and their description is as the following:

#### 1- 2D MALDI FT-ICR MSI of a human prostate dataset:

The analysis was performed according to Randall *et al*[1]. Briefly, 12 µm thickness prostate samples diagnosed with a Gleason score of (3+4)=7 were mounted on a microscopy glass slide and coated with CHCA (5 mg/mL in 70/30 methanol/water with 0.1% trifluoroacetic acid V/V) using an automated sprayer (TM-Sprayer, HTX Imaging, Carrboro, NC). The analysis of the samples was performed on a 9.4 Tesla SolariX XR FT ICR mass spectrometer (Bruker Daltonics, Billerica, MA) using the MALDI source in positive ion mode in the mass range between 250-1000 *m/z*, with a spatial resolution of 120 µm. Internal online calibration was performed using heme *m/z* 616.1776 during data acquisition.

#### 2- 3D MALDI FT-ICR MSI dataset of a PDX mouse brain model of glioblastoma:

The intracranial tumor belonging to a PDX model of GBM12 (PDX National Resource, Mayo Clinic), was analyzed by MALDI FT ICR MSI using a 9.4 Tesla SolariX mass spectrometer (Bruker Daltonics, Billerica, MA), using continuous accumulation of selected ions in the mass range between 380-620 *m/z*. The indium tin oxide (ITO)-coated slide with 12 µm thickness tissue sections , was coated with DHB (160 mg/mL in a 70/30 v/v solution of methanol/0.2% TFA), according to Randall *et al*[2]. The 3D MSI dataset was collected from 4 tissue sections with an inter-slice distance of 160 µm. Internal online calibration was performed using heme *m/z* 616.1776 during data acquisition.

#### 3- 3D DESI MSI dataset of human colorectal adenocarcinoma:

A clinical volumetric tissue specimen of colorectal adenocarcinoma was sliced into 26 sections each of 10 µm thickness and imaged by DESI MSI (Thermo Exactive instrument, Thermo Scientific GmbH, Germany). This 3D MALDI MSI dataset was acquired in the negative-ion mode and covered a mass range of *m/z* 200-1050. The acquisition spatial resolution was set to 100 µm, and the full 3D MSI dataset encompasses 148044 spectra each of 8073 dimensions. This 3D MSI dataset is publicly available and for more information on sample preparations and acquisition details we refer to Oetjen *et al*[3].

#### 4- 3D MALDI MSI dataset of human oral squamous cell carcinoma:

A volumetric tissue specimen of human oral squamous cell carcinoma was sliced into 58 consecutive sections each of 10 µm thickness and imaged by MALDI MSI (Autoflex speed™, Bruker Daltonics, Germany). This 3D MALDI MSI dataset was acquired in the positive-ion mode and covered a mass range of *m/z* 2,000-20,000. The acquisition spatial resolution was set to 60 µm, and the full 3D MSI dataset encompasses 825558 spectra each of 7665 dimensions. This 3D MSI dataset is publicly available and for more information on sample preparations and acquisition details we refer to Oetjen *et al*[3].

#### 5- 3D MALDI MSI data of mouse kidney:

A volumetric tissue specimen of mouse kidney was sliced into 73 consecutive sections each of 3.5 µm thickness and imaged by MALDI MSI (Autoflex speed™, Bruker Daltonics, Germany). This 3D MALDI MSI dataset was acquired in the positive-ion mode and covered a mass range of *m/z* 2,000-20,000. The acquisition spatial resolution was set to 50 µm, and the full 3D MSI dataset encompasses 1,362,830 spectra each of 7671 dimensions. This 3D MSI dataset is publicly available and for more information on sample preparations and acquisition details we refer to Oetjen *et al*[3].

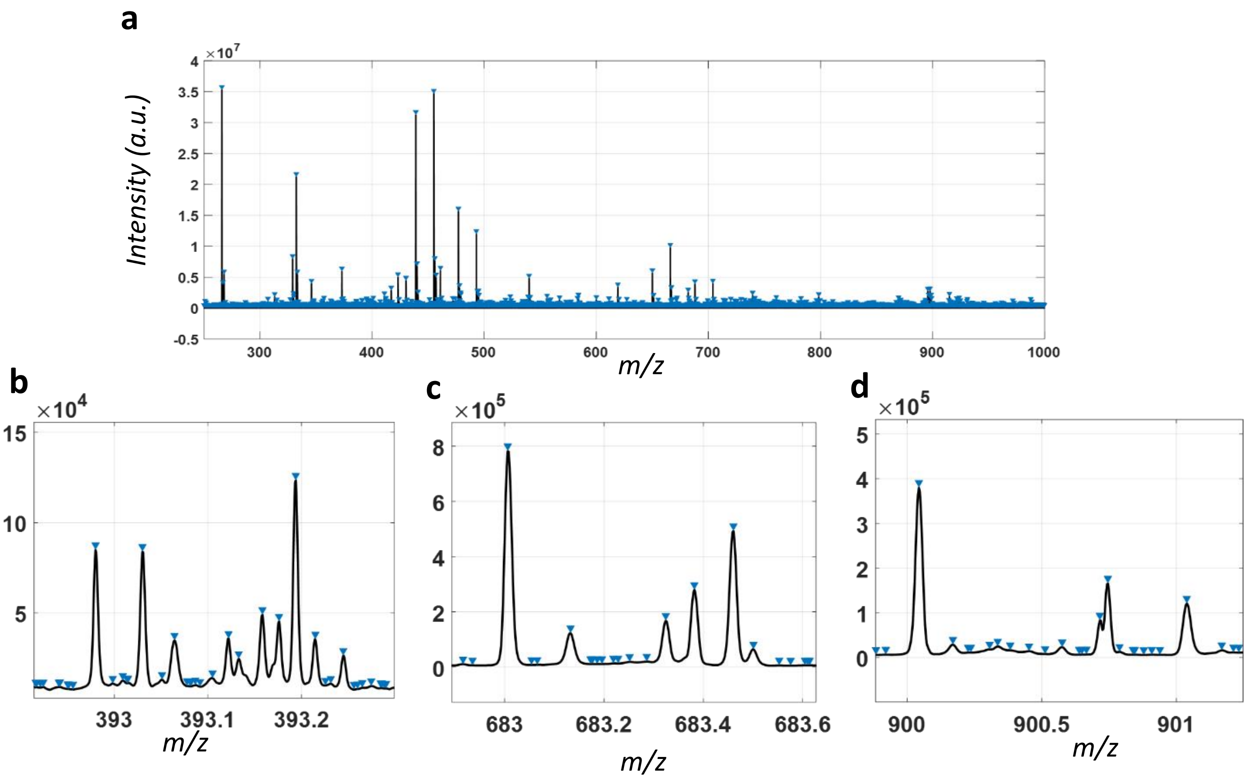


*Figure S1. FT-ICR MSI prostate cancer dataset: local maxima are highlighted in the mean spectrum(a), and a zoomed-in regions within the mean spectrum (b-d). The identified local maxima significantly reduce the original spectral dimensions from about 0.73 million to 61,343 m/z features. This impacts the spectral sparsity but not the spectral representation.*

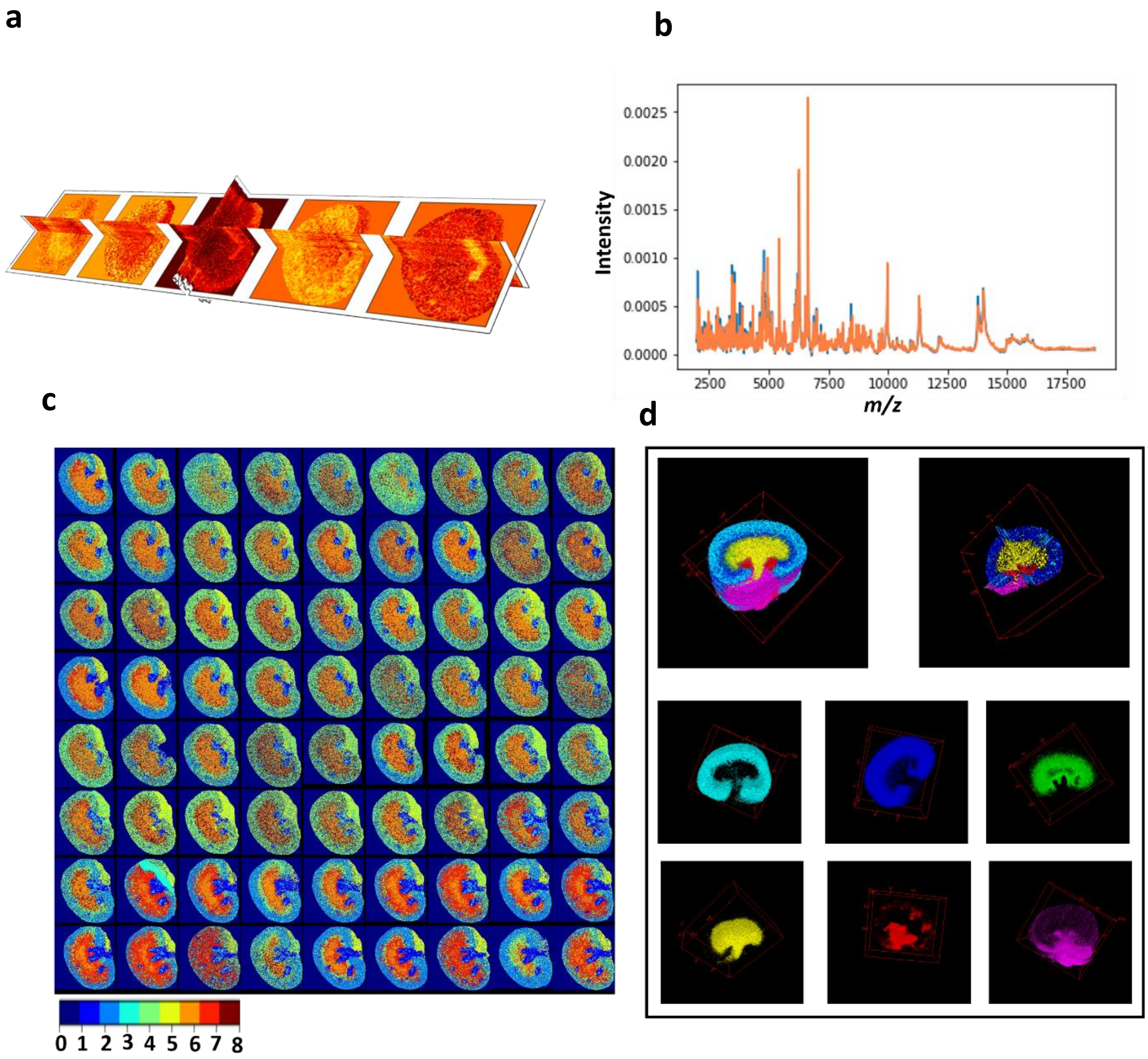


*Figure S2. 3D MALDI MSI dataset from 72 mouse kidney tissue samples that were withheld for test analysis using the trained model shown in the main manuscript Figure 5: a. 3D distribution of the encoded features, b. overlay of the overall mean spectrum of both TIC normalized original (blue) and reconstructed (orange) full test dataset with mean squared error of* $3.11 \times 10^{-3}$*.* ***c****. clustering of encoded features using GMM (k=8) reveals molecular patterns that form anatomical structures that are volumetric rendered in (d).*

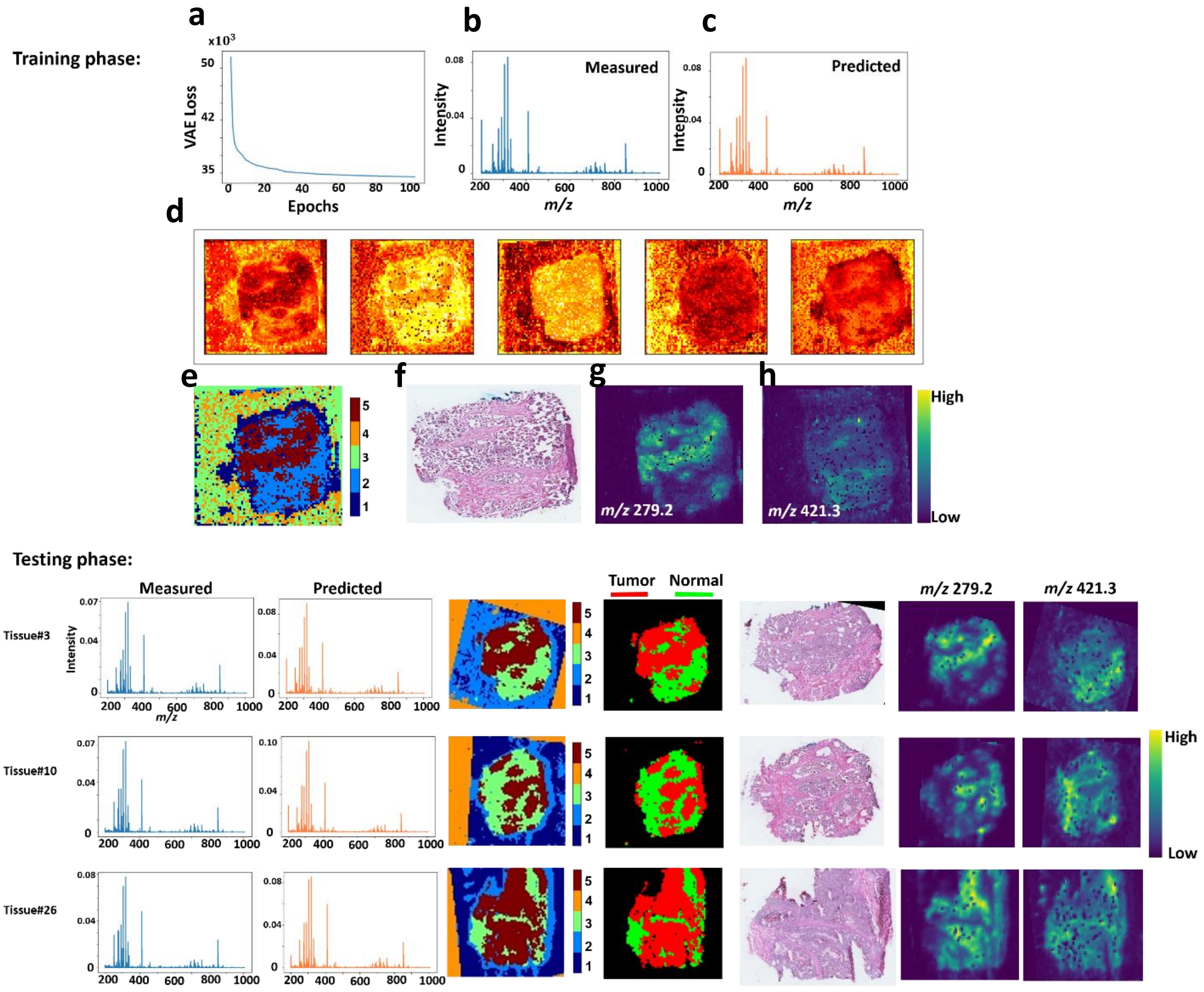


*Figure S3. Deep-Learning based analysis of a 3D DESI MSI dataset of a clinical specimen of colorectal carcinoma. In the* ***training phase*** *the VAE model took around 3.2 minutes to run on a training sample of 5694 spectra each of 8073 dimensions:* ***a.*** *distribution of the optimization convergence with the number of iterations (epochs), average TIC-normalized spectra of both measured (b) and predicted (c) spectral data are closely distributed, d. captured encoded features that visualizes a non-linear embedding of original data, e. these encoded features were clustered with Gaussian mixture model (with k=5) and it revealed two biologically-relevant tissue types that reconcile with the H&E histology (***f***), namely: tumor region (cluster#5) and connective tissue (cluster#2), and distribution of ion features at m/z 279.23 ± 0.1 (***g***) and m/z 421.31 ± 0.1 (***h***) were found elevated and highly correlated with the tumor and connective tissue clusters, respectively. In the* ***testing phase*** *the trained model was applied on the withheld test dataset: results on three distinct test MSI datasets sampled from different locations within the volumetric tissue specimen, each sample was analyzed within 4 seconds in which the model accurately predicted the measured data (overall mean squared error of* $1.77 \times 10^{-4}$*), and molecular-based tumor and connective tissue clusters were identified and found in close agreement with the H&E histology.*

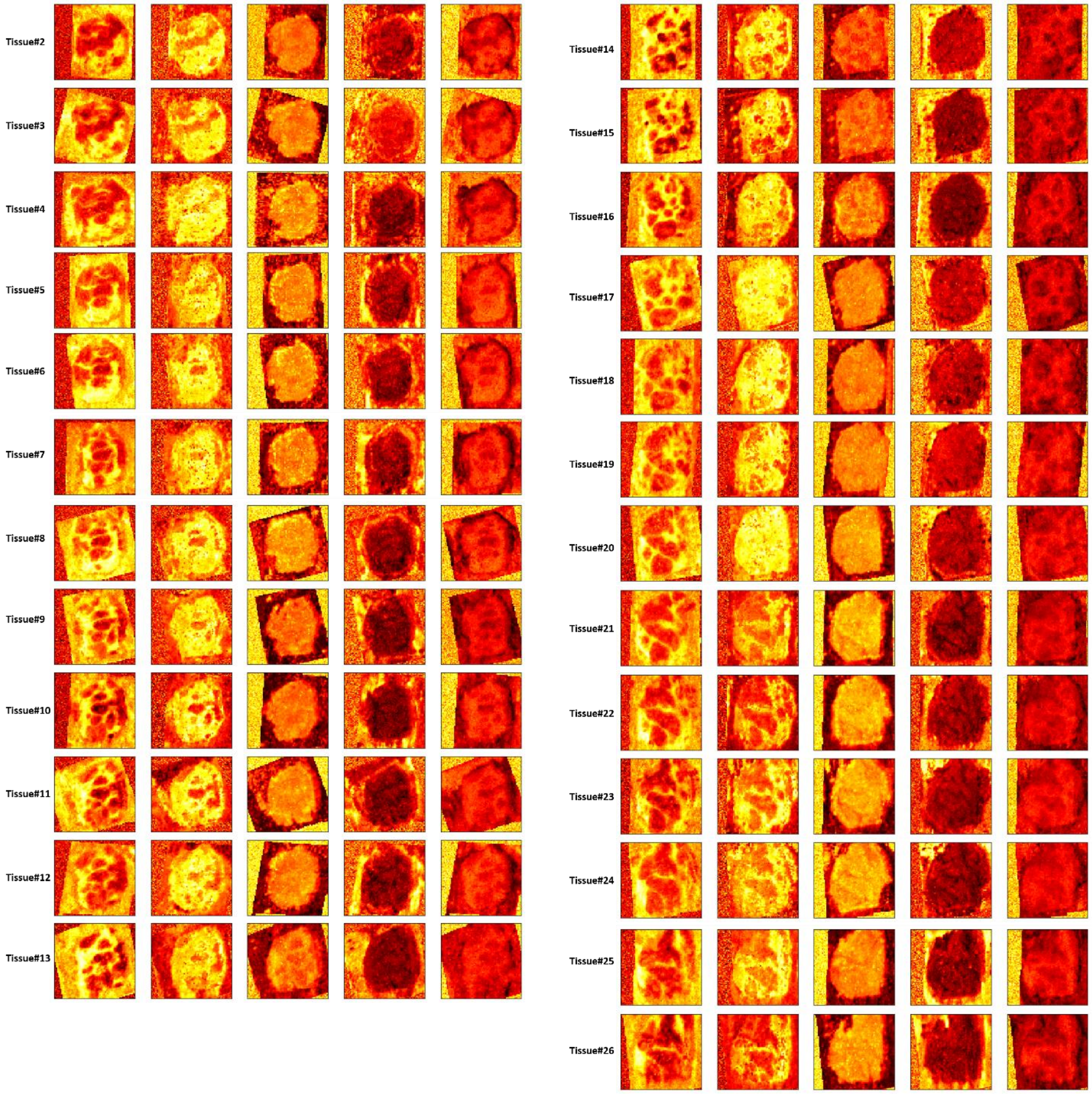


*Figure S4. Encoded features of 3D DESI MSI test dataset of 25 tissue sections of colorectal adenocarcinoma. High dimensional spectral data of each tissue section were non-linearly mapped into a smaller subspace represented by 5 encoded features. The encoded features reveal structural information that depict molecular patterns from original high dimensional data.*

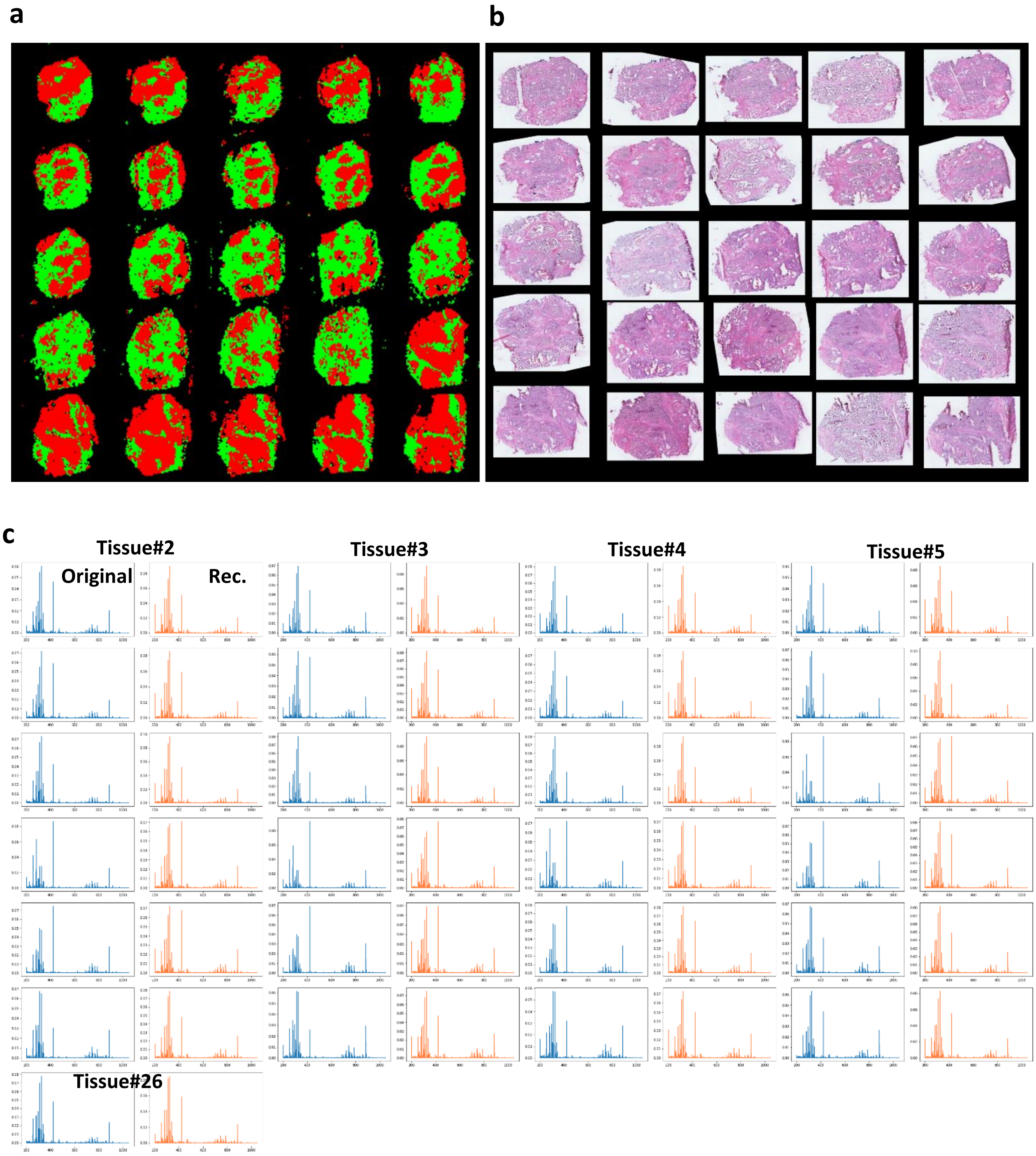


*Figure S5. Analysis of 3D DESI MSI test dataset of colorectal adenocarcinoma: **a.** Tumor (red) and connective tissue (green) clusters were extracted from the clustered image of Gaussian mixture model (k=5) on the encoded features shown in Figure S4, and they appear in a close agreement with their counterparts H&E histology. c. Distribution of average spectrum for both TIC normalized original (blue) and reconstructed (orange) MSI data.*

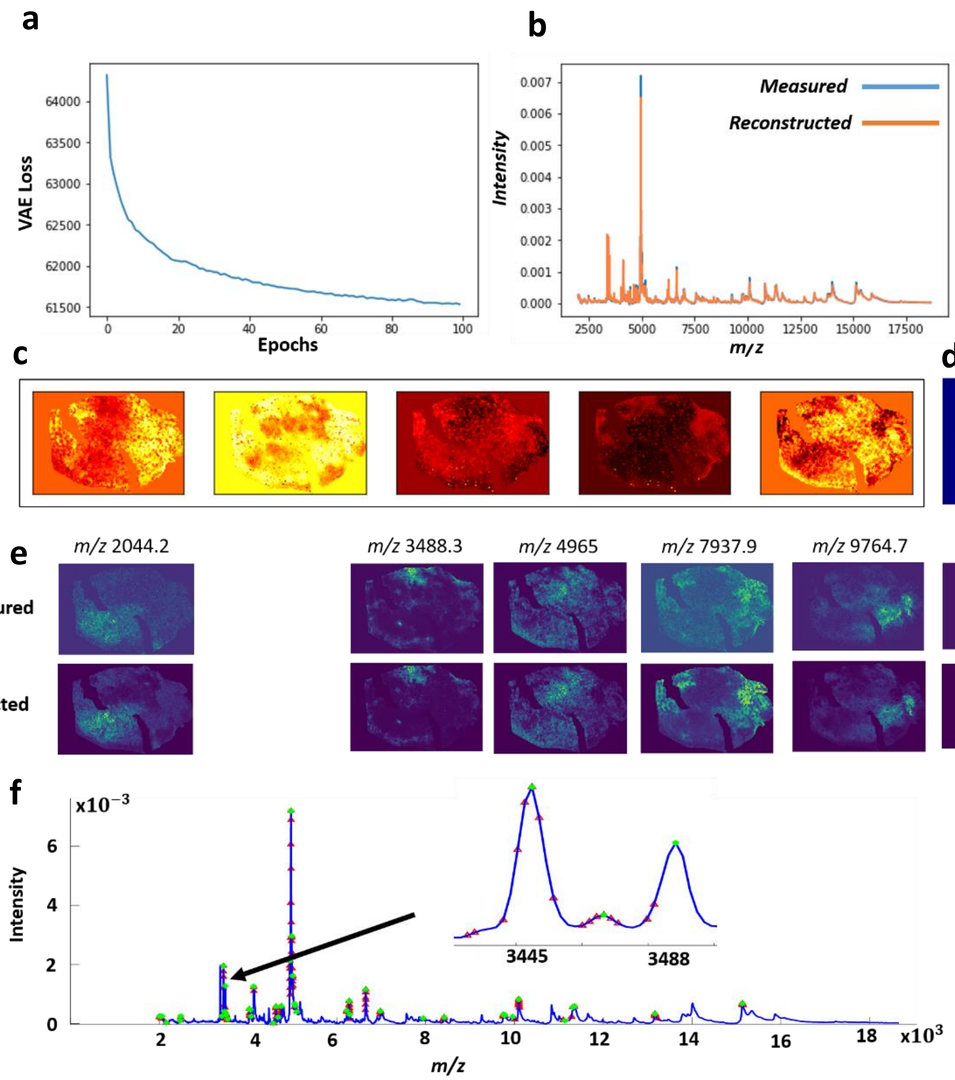


*Figure S6. Deep learning analysis of a 2D MALDI MSI dataset of OSCC tissue:* ***a.*** *Optimizer convergence,* ***b.*** *Overlay of the mean spectrum distribution of TIC normalized original (blue) and reconstructed (orange) MSI data.* ***c.*** *Encoded features of 5 dimensions capture molecular patterns from original high dimensional data and were clustered by the GMM (k=7) to reveal clusters of molecularly distinct regions (****d****). Each of these clusters was correlated with the reduced MSI data of selected peaks (green points in panel* ***f****), and the distribution of highly correlated m/z peaks in each cluster is given in (****d****). The spatial distributions of m/z images (****d****) confirm high quality of reconstructed data. f. High-weighted unbanned m/z variables underlying the molecular structures captured by the encoded features are highlighted in the mean spectrum: m/z bins (red points) and their parent peaks (green points).*

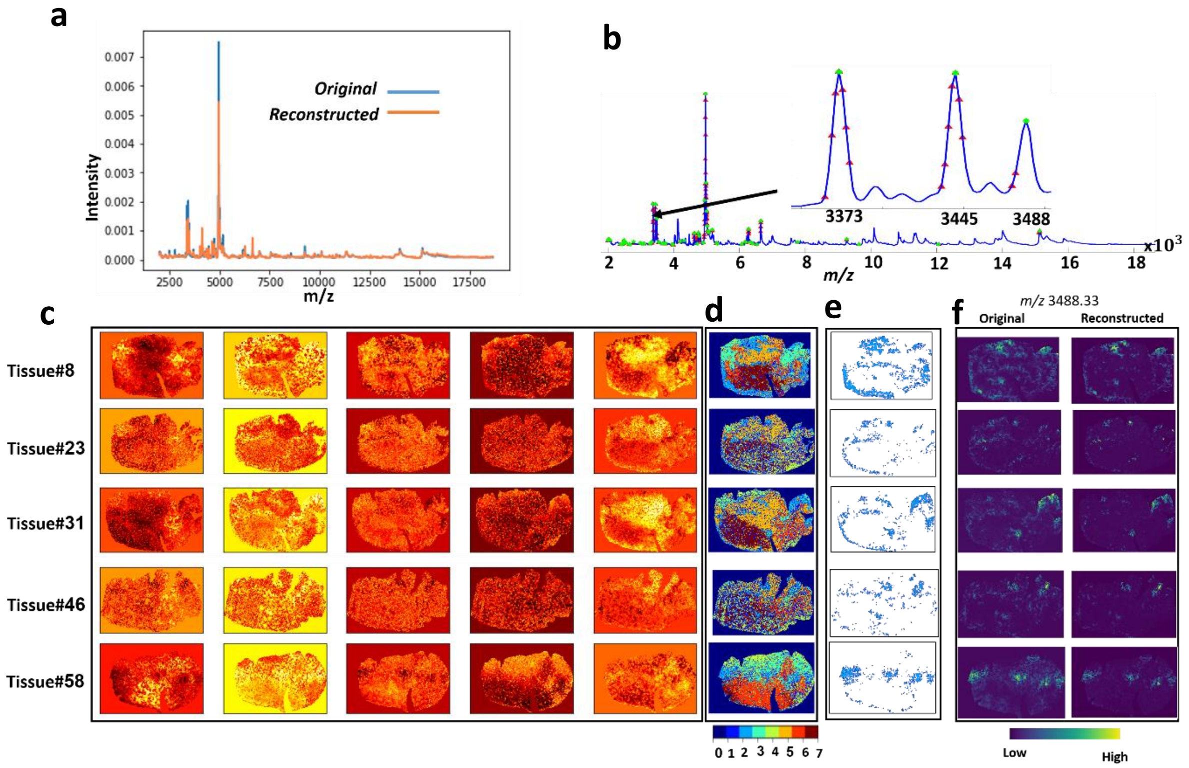


*Figure S7. Analysis of test MSI dataset from 5 tissue sections that were sampled from different locations within the volumetric tissue specimen: a. Overlay of original and estimated TIC normalized mean spectrum, and b. highlights m/z variables (red points highlight m/z bins whereas green points highlight m/z peaks) underlying the molecular patterns depicted in the latent space (c). These molecular patterns were clustered by the GMM (d) and the cluster#2 was extracted (e) and found highly correlated with the defensins produced by the Neutrophils, namely lipid ions at m/z 3373, 3445, and 3488. One of these ions was spatially mapped (f) for both original and reconstructed data.*

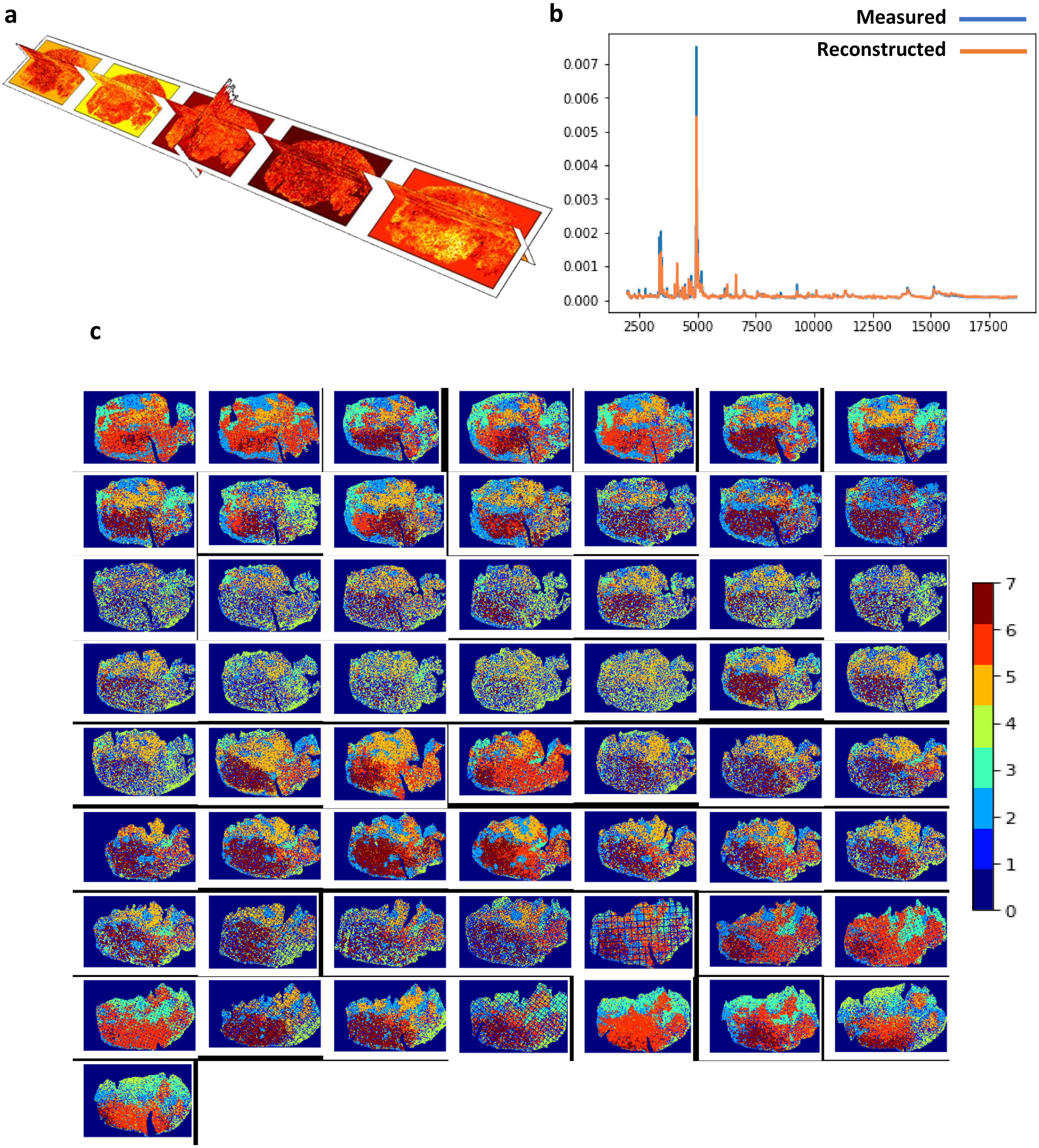


*Figure S8. 3D MALDI MSI dataset from 57 human OSCC tissue samples that were withheld for test analysis using the trained model shown in Figure S7: **a**. 3D distribution of the encoded features, **b**. overlay of the overall mean spectrum of both TIC normalized original (blue) and reconstructed (orange) full test dataset with mean squared error of* $3.03 \times 10^{-3}$. ***c**. clustering of encoded features using GMM (k=8) reveals molecular patterns (for example cluster#2 represents molecular phenotypes of defensins that are produced by the Neutrophils).*

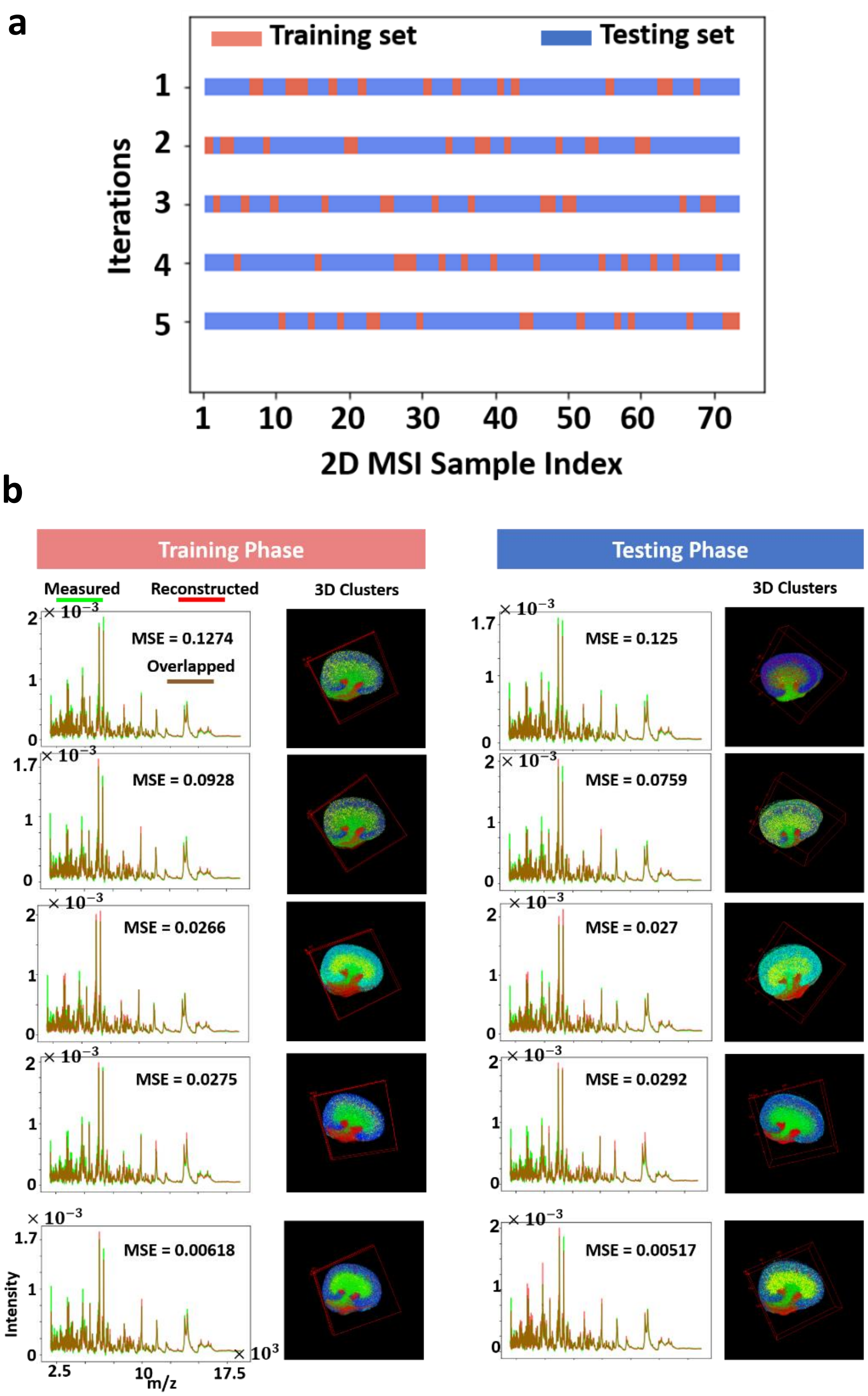


*Figure S9. Cross-validation analysis for the 3D MALDI MSI dataset of the mouse kidney (73 consecutive tissue sections): a. the full MSI dataset was randomly shuffled and split into a 20% training set and an 80% testing set, and this process was repeated 5 times. (b) For each iteration in the cross-validation (each row), the msiPL model was applied on the training set to optimize the artificial neural network and the trained model was then applied on the unseen test set. The original MSI data was completely reconstructed with a small mean squared error (MSE), and the overlay of the average spectrum of both TIC-normalized original and reconstructed data reveals close distribution. The learned non-linear manifold (encoded features) was clustered using GMM(k=8) which revealed distinct molecular patterns that reconcile with the kidney's anatomy. The trained model was robust and did not overfit the test set as it showed comparable performance to the training phase.*

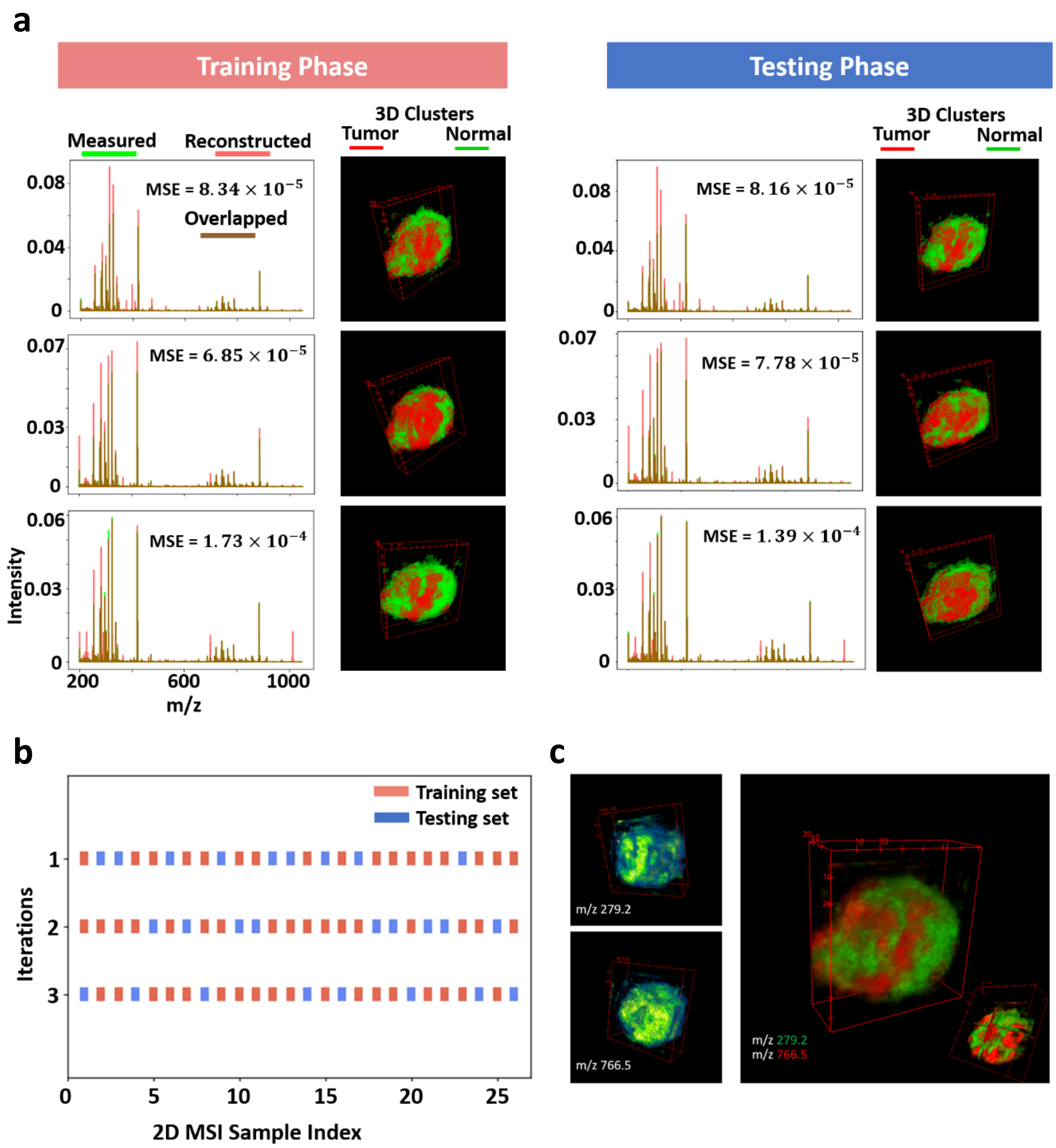


*Figure S10. Cross-validation analysis for the 3D DESI MSI dataset of a human specimen of colorectal carcinoma: (a) the training set was used to optimize the neural network and then the trained model was applied on the testing set, and this process was repeated three times (rows) according to 3-fold cross-validation shown in (b) in which the full dataset was randomly shuffled and split into training and testing sets. There is a close consensus in the performance of the cross-validated models in reconstructing the original data, learning the non-linear manifold and identifying the tumor and normal clusters. (c) The three cross validated models showed stability in learning peaks of interest such as m/z 279.2 and m/z 766.5 that were found localized (>0.7 Pearson correlation) and elevated in the tumor and normal clusters respectively.*

(a) Using the commercial software:

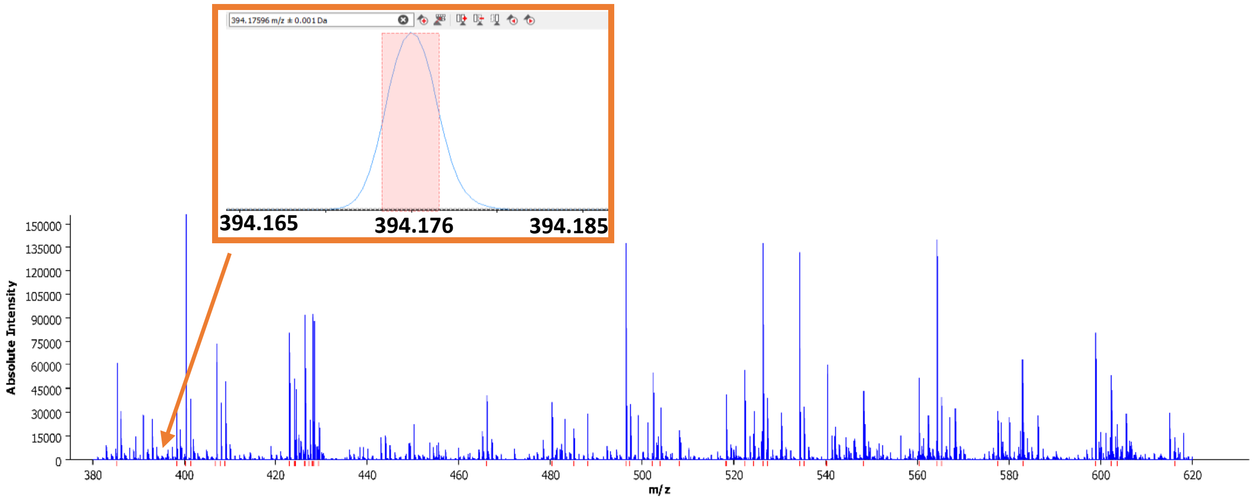


(b) Using msiPL:

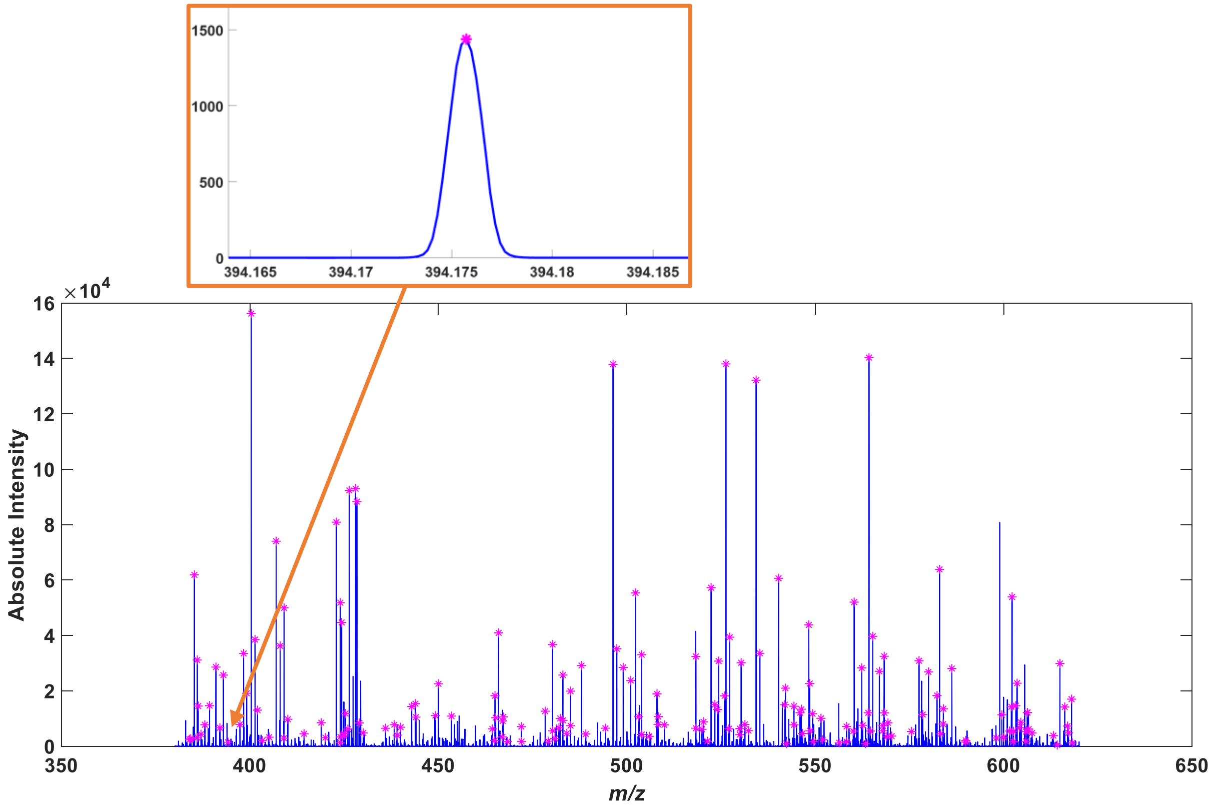


*Figure S11. Peaking picking on the 3D MSI dataset of PDX mouse brain tumor of glioblastoma using the default parameters of the commercial software SCiLS 2019c (Bruker, Germany) and the proposed msiPL method: (a) The identified peaks are highlighted with red points on the average spectrum using the commercial software. The commercial software default parameters are global, and it could not identify an important peak corresponding to the erlotinib drug metabolite at m/z* $394.175 \pm 0.001$ *which is present in the data as shown in the inset. (b) The learned peaks using the msiPL method are highlighted in the average spectrum and revealed that the erlotinib drug metabolite at m/z 394.1757 is identified unlike the commercial software with default parameters.*

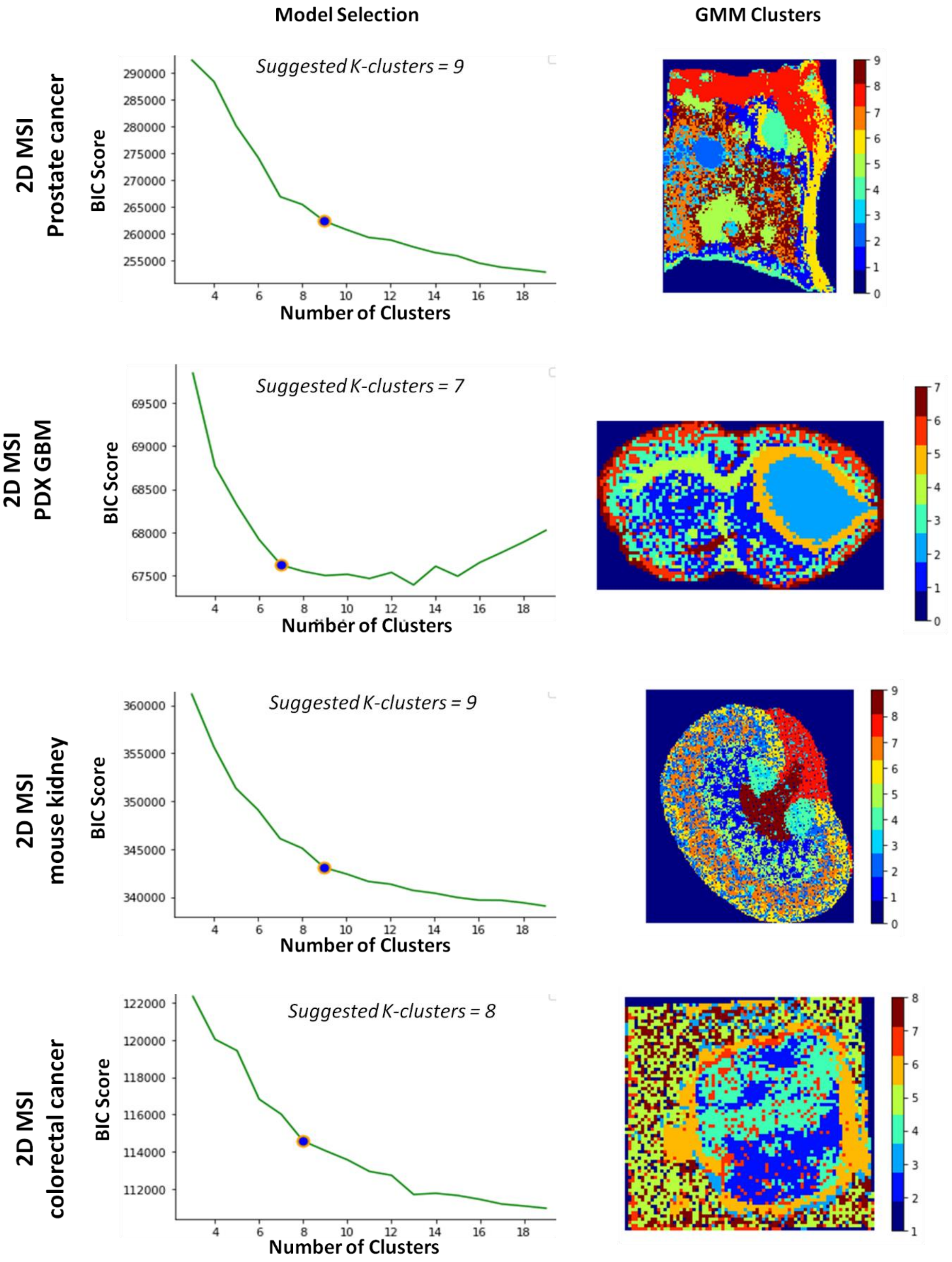


*Figure S12.* ***Optimization-based selection of the number of K-clusters:*** *Model selection of K-clusters using Bayesian information criterion (BIC) and the Kneedle algorithm to identify the optimum model at the point of maximum curvature (first column), and the spatial distribution of GMM clusters based on the suggested number clusters (second column)..*

***Table S1.*** *FT-ICR MSI prostate dataset: highly correlated m/z ion peaks with the prostate molecular-based tumor cluster*

| *m/z* experimental | correlation | *m/z* Identity | Molecular formula | Adduct | *m/z* calculated | Database | Error (ppm) |
|---|---|---|---|---|---|---|---|
| 739.4664 | 0.7015 | PA | C39H73O8P | [M+K] | 739.4675 | HMDB | 1.44 |
| 763.4662 | 0.6852 | PA | C41H73O8P | [M+K] | 763.4675 | HMDB | 1.66 |
| 737.4506 | 0.6669 | PA | C39H71O8P | [M+K] | 737.4518 | HMDB | 1.65 |
| 985.5567 | 0.6519 | PIP(P-42:6) | C51H88O15P2 | [M+H-H2O]+ | 985.5566 | LipidMaps | -0.14 |
| 761.5867 | 0.6472 | | | | | | |
| 760.5827 | 0.6462 | | | | | | |
| 738.4548 | 0.6462 | PI-Cer(t30:2) | C36H68NO12P | [M+H]+ | 738.4552 | LipidMaps | 0.53 |
| 786.5981 | 0.6441 | PC(O-34:0(OH)) | C42H86NO8P | [M+Na]+ | 786.5983 | LipidMaps | 0.29 |
| | | PE(O-37:0(OH)) | C42H86NO8P | [M+Na]+ | 786.5983 | LipidMaps | 0.29 |

***Table S2.*** *Highly correlated m/z ion peaks with the tumor rim (Cluster#4) of the FT-ICR MSI PDX mouse brain dataset*

| m/z experimental | correlation | *m/z* Identity | Molecular formula | Adduct | *m/z* calculated | Database | Error (ppm) |
|---|---|---|---|---|---|---|---|
| 438.2979 | 0.515 | Palmitoylcarnitine | $C_{23}H_{45}NO_4$ | [M+K]+ | 438.2980 | HMDB | 0.27 |
| 464.3132 | 0.484 | Carnitine | $C_{25}H_{47}NO_4$ | [M+K]+ | 464.3137 | HMDB | 1.01 |
| 400.3419 | 0.464 | Palmitoylcarnitine | $C_{23}H_{45}NO_4$ | [M+H]+ | 400.3421 | HMDB | 0.59 |
| 428.3734 | 0.463 | Stearoylcarnitine | $C_{25}H_{49}NO_4$ | [M+H]+ | 428.3734 | HMDB | 0.08 |
| 401.3454 | 0.459 | | | | | | |
| 398.3259 | 0.458 | 9-Hexadecenoylcarnitine | $C_{23}H_{43}NO_4$ | [M+H]+ | 398.3265 | HMDB | 1.47 |

***Table S3.*** *Highly correlated m/z ion peaks with the tumor region1 (Cluster#2) of the FT-ICR MSI PDX mouse brain dataset*

| *m/z* experimental | correlation | *m/z* Identity | Molecular formula | Adduct | *m/z* calculated | Database | Error (ppm) |
|---|---|---|---|---|---|---|---|
| 567.9409 | 0.733 | | | | | | |
| 605.8966 | 0.73 | | | | | | |
| 589.923 | 0.728 | | | | | | |
| 529.9846 | 0.718 | ATP/dGTP | C10H16N5O13P3 | [M+Na]+ | 529.9850 | HMDB | 0.69 |
| 589.9102 | 0.711 | | | | | | |
| 545.959 | 0.707 | ATP/dGTP | C10H16N5O13P3 | [M+K]+ | 545.9589 | HMDB | -0.18 |
| 464.9473 | 0.704 | | | | | | |
| 567.9285 | 0.701 | | | | | | |
| 583.9147 | 0.7 | | | | | | |
| 605.8842 | 0.699 | | | | | | |

***Table S4.*** *Highly correlated m/z ion peaks with the tumor region2 (Cluster#8) of the FT-ICR MSI PDX mouse brain dataset*

| *m/z* experimental | correlation | *m/z* Identity | Molecular formula | Adduct | *m/z* calculated | Database | Error (ppm) |
|---|---|---|---|---|---|---|---|
| 558.2953 | 0.743 | LysoPC(18:2) | C26H50NO7P | [M+K]+ | 558.2956 | HMDB | 0.62 |
| 520.3397 | 0.737 | LysoPC(18:2) | C26H50NO7P | [M+H]+ | 520.339766 | HMDB | 0.13 |
| 468.3083 | 0.729 | LysoPC(14:0) / LysoPE | C22H46NO7P | [M+H]+ | 468.3084 | HMDB | 0.35 |
| 414.3579 | 0.727 | Heptadecanoyl carnitine | C24H47NO4 | [M+H]+ | 414.3577 | HMDB | -0.28 |
| 521.3428 | 0.717 | | | | | | |
| 425.3453 | 0.716 | | | | | | |
| 424.3421 | 0.713 | Linoelaidyl carnitine or Linoleyl carnitine | C25H45NO4 | [M+H]+ | 424.3421 | HMDB | 0.08 |
| 494.3239 | 0.709 | Cervonyl carnitine | C29H45NO4 | [M+Na]+ | 494.3240 | HMDB | 0.36 |
| | | LysoPC(16:1) | C24H48NO7P | [M+H]+ | 494.3241 | HMDB | 0.44 |
| 532.28 | 0.7 | LysoPC(16:1) | C24H48NO7P | [M+K]+ | 532.2799 | HMDB | 0.00 |
| 426.3573 | 0.697 | Carnitine | C25H47NO4 | [M+H]+ | 426.3577 | HMDB | 1.14 |

***Table S5.*** *Different parameter settings of the neural network model (tested on a 2D MALDI MSI data of PDX mouse brain model of glioblastoma)*

| Design | #Hidden Layers | #Computational Neurons per Layer | #Model Parameters | Running Time (Minutes) | MSE of Original and Reconstructed Data |
|---|---|---|---|---|---|
| Architecture1 | 3 Layers | 50-5-50 | 2,146,421 | 3.1 | $6.52 \times 10^{-4}$ |
| Architecture2 | 3 Layers | 512-3-512 | 21,779,723 | 3.2 | $4.72 \times 10^{-4}$ |
| Architecture3 | 3 Layers | 512-5-512 | 21,782,807 | 3.6 | $4.56 \times 10^{-4}$ |
| Architecture4 | 3 Layers | 512-15-512 | 21,798,227 | 3.6 | $3.47 \times 10^{-4}$ |
| Architecture5 | 3 Layers | 5000-5-512 | 212,536,271 | 8.27 | $2.82 \times 10^{-4}$ |
| Architecture6 | 5 Layers | 3000-512-5-512-3000 | 120,568,023 | 5.8 | $2.02 \times 10^{-4}$ |
| Architecture7 | 7 Layers | 5000-1000-512-5-512-1000-5000 | 232,502,023 | 8.4 | $2.28 \times 10^{-4}$ |

***Table S6.*** *Running time of training the msiPL on the GPU compared to the CPU.*

| Dataset | msiPL on CPU | msiPL on GPU |
|---|---|---|
| 2D FT ICR MSI of human prostate cancer (12,716 spectra; 730,403 m/z bins) | 149.8 minutes | 40 minutes |
| 2D MSI of mouse kidney (18,536 spectra; 7671 m/z variables) | 25.2 minutes | 8.6 minutes |

***Table S7.*** *Running Time for peak picking using commercial software compared to msiPL*

| Dataset | Running Time: Only Peak Picking Using SCiLS 2019c (Bruker, Germany) | The entire msiPL analysis (not just the peak learning step) |
|---|---|---|
| 2D FT ICR MSI of human prostate cancer (12,716 spectra; 730,403 m/z bins) *<br>*sampling every 20th spectrum | 4 hours and 50 minutes | 40 minutes |
| 3D MALDI MSI of a PDX mouse brain model of glioblastoma (14,833 spectra; 661,402 m/z bins)+<br>+sampling every 5th spectrum | 10 hours and 39 minutes | Training phase: 3.6 minutes Testing phase: 8 seconds.<br><br>Note: 3D MSI dataset is analyzed using msiPL using training/testing strategy as explained in the manuscript. |
